# Co-recruitment of relatives in a broadcast-spawning coral (*Acropora hyacinthus*) facilitates emergence of an inbred, genetically distinct group within a panmictic population

**DOI:** 10.1101/2020.02.26.956680

**Authors:** Sarah Barfield, Sarah W. Davies, Mikhail V. Matz

## Abstract

Many broadly-dispersing marine taxa are species-rich, show genetic heterogeneity on small spatial scales, and are locally adapted to their environmental conditions. How such genetic subdivisions can emerge despite the potential for high gene flow continues to be the major paradox of evolution in the sea. One understudied process potentially contributing to genetic structuring in marine populations is variation in larval recruitment. Here, we report an unusual recruitment pattern in the broadcast-spawning coral species *Acropora hyacinthus* on Yap Island, Micronesia. Reduced representation genotyping of 275 individuals of varying size classes on this isolated reef system demonstrated general island-wide panmixia but also identified a genetically divergent group of juvenile corals at one out of the four sites sampled, showing elevated inbreeding and relatedness, including two pairs of siblings. Notably, adult corals as well as the majority of juveniles at the same site belong to the panmictic gene pool, suggesting that representatives of the inbred lineage of juveniles co-recruited and may be partially reproductively isolated from the rest of the island population. Reproductive isolation is suggested by the discovery of distinct genomic regions of greatly reduced genetic diversity in the inbred lineage, encompassing genes involved in sperm-egg recognition and fertilization that may serve as reproductive barrier loci. We propose that co-recruitment of genetic relatives via cohesive dispersal, a process that was previously unrecognized in broadcast-spawning corals, can generate familial genetic structure and might be important for the emergence of genetically distinct, locally adapted ecomorphs and cryptic species.

## Introduction

Broadcast spawning marine invertebrates, such as corals, often have very large population sizes, produce planktonic larvae with long pelagic durations, and exhibit low genetic differentiation over large geographic ranges (Ayre & Hughes, 2000; Ayre & Hughes, 2004; Davies, Treml, Kenkel, & Matz, 2015; Van Oppen, Peplow, Kininmonth, & Berkelmans, 2011). The seemingly extensive gene flow that characterizes broadcast spawning marine organisms poses a conundrum, since high levels of gene flow can slow or inhibit local adaptation, especially if migration swamps a local population with maladaptive genetic variation (Garant, Ford, & Hendry, 2007; Lenormand, 2002). Yet, many marine invertebrate taxa are very species rich, such as the coral genus *Acropora*, despite these species having extensive, overlapping ranges (Palumbi, 1994; Rex et al., 2000; Van Oppen, McDonald, Willis, & Miller, 2000; Wallace & Rosen, 2006; Wilson & Hessler, 1987). This leads to the longstanding question of how genetic subdivisions evolve among marine populations with high dispersal potential. If selection is strong enough along an environmental gradient, this could produce a barrier to genetic exchange among loci underlying adaptation (DeFaveri, Jonsson & Merila, 2013; Meester, Gomez, Okamura, & Schwenk, 2002); but such selection must be incredibly strong to counteract the homogenizing effect that migration would have on the rest of the genome (Mullen & Hoekstra, 2008). Furthermore, it is uncertain how sympatric marine invertebrate populations might make the leap from a locally adapted deme to a reproductively isolated species, since reproductive isolation would lead to a very low chance of fertilization outside of a small deme of reproductively compatible individuals (Munday, Van Herwerden, & Dudgeon, 2004; Palumbi,1994).

The majority of corals species that are critical constituents in the formation of reefs employ a broadcast spawning reproductive strategy. The coral genus *Acropora* in particular is both species rich, having undergone rapid diversification in the past 1-2 million years (Simpson, Kiessling, Mewis, Baron-Szabo, & Muller, 2011; Van Oppen, McDonald, Willis, & Miller, 2000), and an ideal system with which to address questions of how genetic subdivisions can evolve in marine species with planktonic larvae. Some acroporid species, including *Acropora hyacinthus*, are actually comprised of multiple sympatric cryptic species, yet it is not well understood how such species form (Ladner & Palumbi, 2012). Furthermore, understanding the processes by which genetic subpopulations and/or locally adapted populations of corals can develop despite the potential for high migration from other sources is relevant for forecasting coral population trajectories as they adapt to global warming (Matz, Treml, Aglyamova, & Bay, 2018).

Larval recruitment is one process that is relevant for understanding the emergence of genetic subdivisions, since fertilization of egg and sperm generally occurs between closely situated corals on the reef (Babcock, Mundy & Whitehead, 1994; Levitan & Petersen, 1995; Yund, 2000), and thus the extent of self- or co-recruitment may dramatically affect the overall level of outbreeding in a population. Here, we examined genetic signatures across age cohorts of *A. hyacinthus* on Yap Island, Micronesia to investigate the potential for spatial and temporal variation in recruitment dynamics to influence genetic structure in the broader population. We have extensively sampled adult and juvenile *A. hyacinthus* cohorts at four separate locations on Yap Island in order to infer genetic sources of newly established juveniles within these locations and compare juvenile genetic structure to adults on the same reef. Based on our results, we discuss the potential for cohesive larval dispersal to facilitate the emergence of fine-scale genetic structure in the sea amidst general panmixia.

## Methods

### Coral Sampling and Transects

In November 2015, adult and juvenile *Acropora hyacinthus* colonies were sampled from four separate sites on Yap Island (South Tip (9° 26’ 3.1992’’ N, 138° 2’ 13.9488’’ E), West Outer Reef (9° 33’ 33.9336’’ N, 138° 5’ 22.1496’’ E), Nimpal (9° 34’ 0.2496’’ N, 138° 5’ 50.1324’’ E), Goofnuw Channel (9° 34’ 16.41’’ N, 138° 12’ 10.3284’’ E)) (Fig. 1A, 1B).

**Figure 1:**
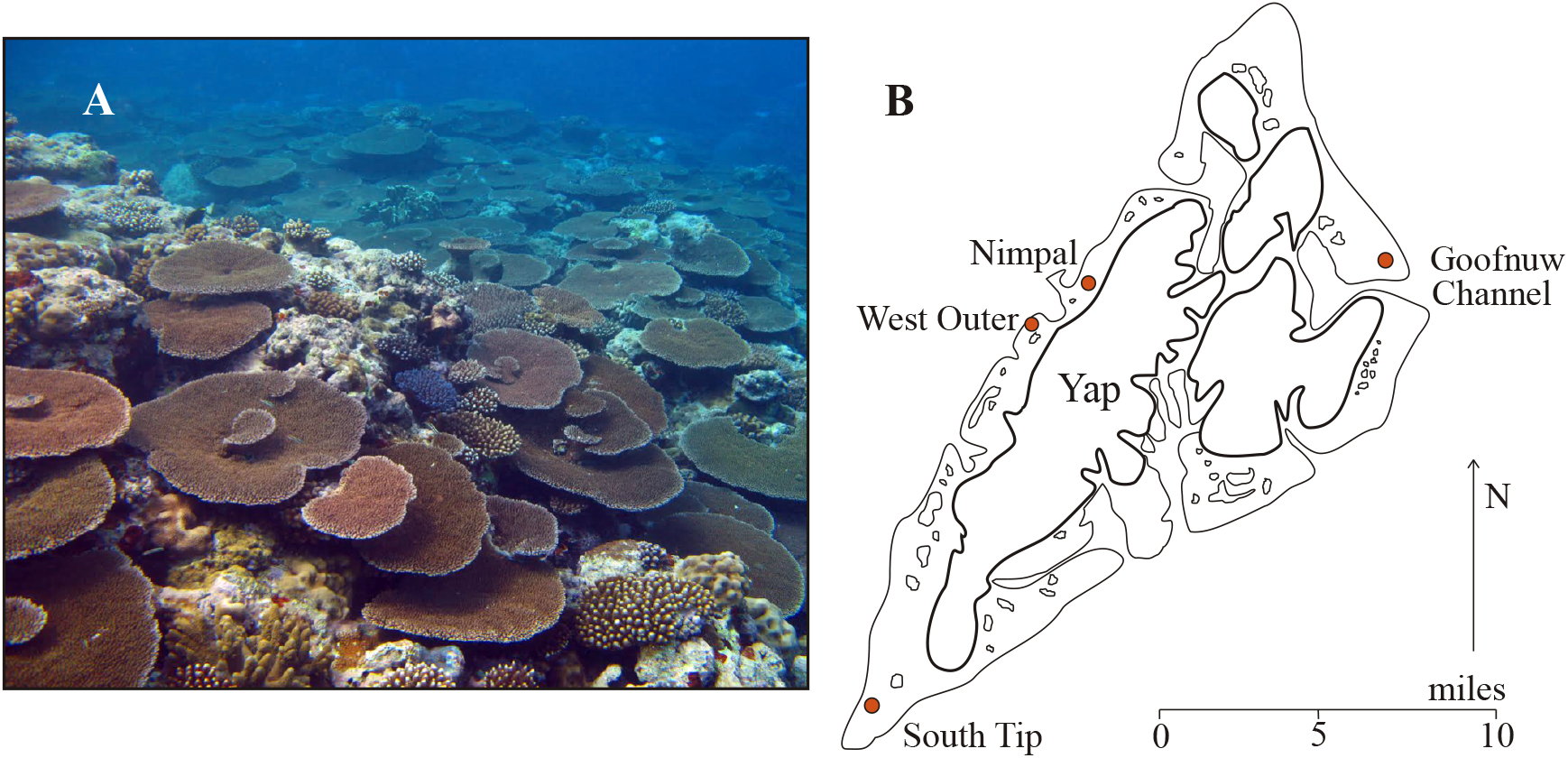
A) Healthy stands of large *A. hyacinthus* present in 2009 at South Tip, Yap Island (Photo Credit: Sarah Davies). B) Sampling locations for *A. hyacinthus* individuals used in this study, including South Tip, West Outer Reef, Nimpal Reef, and Goofnuw Channel.

Only juveniles were collected from South Tip during November 2015 as no adults were present on the reef following the crown-of-thorns starfish outbreak a few years prior. Adult *A. hyacinthus* samples from the South Tip site (2009) were collected by Davies *et al.* a year prior to the outbreak (Davies, Treml, Kenkel, & Matz, 2015). The number of samples per age cohort and location are shown in Supp. table 1.

To sample colonies, small branches 1-2 cm in length were removed from *A. hyacinthus* colonies using pliers and stored in 100% ethanol at −20° C until they could be processed to extract DNA. Photographs with a size standard were taken for each colony in order to determine the sizes of juveniles and adults at each site. Juveniles were classified as individuals with a diameter of less than 20 cm and adults as individuals with a diameter greater than 30 cm. The median diameter and area of juvenile colonies was 8.9 cm and 64.1 cm^2^, respectively (supp. fig. 1), and 60.2 cm and 1,910 cm^2^ for adult colonies (supp. fig. 2).

To determine the composition and relative abundance of adult and juvenile acroporids at each site, three 50 m parallel transects were performed at each site, spaced approximately 5-10 m apart in the vicinity of where coral samples were collected. For each transect, our Yapese field assistants from Yap Community Action Program (Yap CAP) laid out a 50 m reel of transect tape along the reef and recorded video with an underwater camera directly above the tape.

### Sample preparation, sequencing, and image analysis

DNA was extracted from all coral samples using a modified phenol-chloroform extraction method (Davies et al., 2013), and then subsequently genotyped with the 2b-RAD method of Wang *et al*. (Wang, Meyer, Mckay, & Matz, 2012). To ensure accurate genotyping, six samples were replicated three times to identify repeatable sequenced SNPs representing true genetic variation. The final library preparations were combined after barcoding and sequenced on six lanes of the Illumina HiSeq 4000 at UT Austin. Raw sequence data can be found on NCBI-SRA, accession no. PRJNA565239.

To assess the size distributions of sampled corals, photographs were analyzed in ImageJ by using measurements on a size standard placed next to each coral in the photograph. Both diameter and area were calculated for each coral. If the size standard was missing in the photograph, the width of an individual *A. hyacinthus* coral branch (4 mm) was used as the proxy size standard, since this size varies little between individuals.

Analysis of the transect videos was performed by counting the number of adult acroporid colonies, juvenile acroporid colonies, and adult and juvenile colonies of all other coral genera that were in contact with or underneath the transect tape in the video. Since there are some acroporid species on Yap Island that look remarkably similar to *A. hyacinthus* and can be difficult to distinguish at the juvenile stage, we did not try to distinguish *A. hyacinthus* from the other acroporids in video analyses. However, it is worth noting that *A. hyacinthus* is one of the most abundant coral species on Yap Island.

### Reference-based Genotyping

A total of 299 samples were sequenced and genotyped using the *Acropora millepora* genome as a reference. Prior to mapping, reads were trimmed (barcodes removed), deduplicated, and quality filtered with FASTX TOOLKIT such that only reads in which 90% or more of the bases with a Phred score of at least 20 were retained. These reads were subsequently mapped to the newly available *A. millepora* genome (Fuller, Liao, Bay, Matz, & Przeworski, 2019) with bowtie2 (Langmead & Salzberg, 2013). Genotyping was performed with ANGSD v0.921, which utilizes genotype probabilities instead of genotype calls, such that later analyses can take into account genotype uncertainty (Korneliussen, Albrechtsen, & Nielsen, 2014). This characteristic is useful for low or medium coverage data. Only loci with a depth of at least 6 in non-missing individuals were retained for analysis (supp. fig. 3A-C). Additionally, only loci that had a minimum mapping quality of 20, a minimum base call quality of 30, a minor allele frequency of a least 0.05, and were genotyped in at least 80% of individuals were retained, resulting in 6,946 genotyped polymorphic sites.

### Principal coordinates analysis, Admixture, and F_ST_

Principal coordinates analysis (PCoA) utilizing covariance matrices generated by the ANGSD subprogram ngsCovar was performed with the R package “vegan” with the capscale function (constrained analysis of principle coordinates). Samples were partitioned according to sampling location and age cohort. Adults and juveniles from the same location were compared for each PCoA to determine the extent of genetic similarity across age groups.

To further refine patterns of genetic structure among all samples, we used the program NgsAdmix to partition all samples between two genetic clusters (Skotte, Korneliussen, & Albrechtsen, 2013). For clonal pairs identified at Goofnuw Channel, one individual of the pair was removed for the admixture analysis, since clonemates would form their own admixture clusters, obscuring the results. Individuals with genetic variation present in both clusters are shown as bi-colored bars in the admixture plots. Genome-wide F_ST_ estimates were determined with the ANGSD subprogram realSFS. Only weighted F_ST_ estimates are reported to account for differences in sample sizes.

### Population genetic statistics

A hierarchical clustering dendrogram based on the identity-by-state matrix generated by ANGSD was created in R with the *hclust* function using method “average”. Lengths of the branches in the diagram indicate overall genetic similarity among samples, with longer branches shared between two or more samples indicating greater similarity.

Pairwise relatedness, as well as inbreeding coefficients, were calculated with the ANGSD subprogram ngsRelate, which utilizes genotype likelihoods and population allele frequencies. Theta, the measure of genetic diversity (expected number of differences between two homologous sequences randomly chosen from the population), was calculated in windows of 50 kb with step size of 10 kb using the ANGSD subprogram ThetaStat. Heterozygosity was calculated using genotype likelihoods generated by ANGSD and an expectation maximization algorithm (EM) utilized by realSFS in a custom script (heterozygosity_beagle.r), available on the github page associated with this paper (git@github.com:sbarfield/Yap_Ahyacinthus-.git). A Welch two sample t-test was used to identify significant differences in estimates of inbreeding, heterozygosity, and pairwise relatedness.

### Read Depth Analysis

To ensure our analyses were not biased by low-coverage sequencing of some individuals, ^we performed a principal component analysis on the log_10_ of read depth, heterozygosity,^ inbreeding coefficients, and percent blue ancestry. In addition, we compared the distribution of read depths amongst samples, including putative relatives and Nimpal juveniles.

## Results

### Adult and Juvenile Coral Density on Yap Island

Analyses of coral abundances based on transects for both acorporid and non-acroporid corals at all four sampling sites indicate that Goofnuw Channel had the lowest numbers of both adult and juvenile colonies (supp. fig. 4). In addition, owing to the small number of juveniles at this site, they tended to have a larger range of surface area on average than juveniles from the other three sites (supp. fig. 4). In contrast, Nimpal, West Outer, and South Tip reefs had much higher coral cover, and acroporid coral species tended to be more abundant than non-acorporid species. At South Tip, the average number of acroporid adults per transect was 21 colonies compared to 33 colonies for Nimpal and 66 colonies for West Outer Reef. In contrast, the average number of acroporid juveniles per transect at South Tip was much higher than adults. Anecdotally, while sampling corals at South Tip, we noticed very few *A. hyacinthus* adult colonies, in line with our expectation of reduced acroporid adult population sizes due to a prior crown-of-thorns sea star outbreak.

### Genetic Structure of Adult and Juvenile Cohorts on Yap Island

Analysis of principle coordinates (PCoA) based on pairwise genetic distances for all adult and juvenile cohorts demonstrates the lack of genetic structure between sites and also between age cohorts, with the notable exception of Nimpal adults and juveniles, which show separation along the first principle coordinate (fig. 2A-D). South Tip and West Outer Reef juveniles, surprisingly, show nearly complete overlap in principal coordinate space with their respective adult counterparts. This is despite the fact that, at least for South Tip, adults were sampled five years prior and were then lost to the crown-of-thorns outbreak.

**Figure 2:**
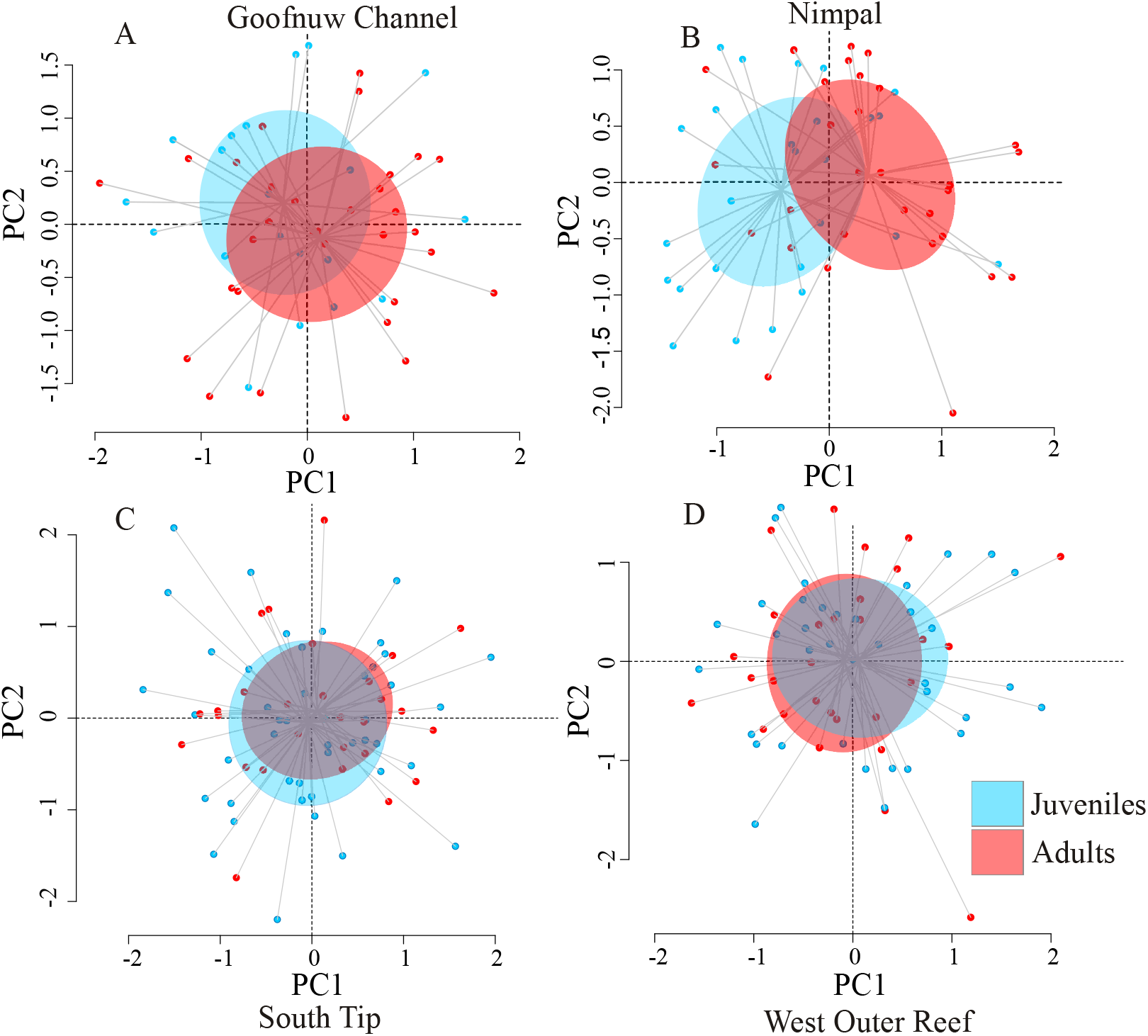
Principal coordinates analysis (PCoA) based on genetic covariance matrices of adult and juvenile population cohorts. Red points represent adult individuals and blue points represent juvenile individuals. A) Goofnuw Channel B) Nimpal Reef C) South Tip D) West Outer Reef.

Further analysis of population structure with ngsAdmix demonstrates no significant separation of genetic variation between sites or cohorts when clustered into two admixture groups, with the exception of Nimpal adults and juveniles, as was demonstrated with PCoA (fig. 3). Nearly half of the Nimpal juveniles show 100% genetic affiliation with a separate population cluster (blue), along with two South Tip adults, one South Tip juvenile, and one West Outer Reef juvenile (fig. 3). All other individuals across sites display a majority red ancestry, but also show at least some level of admixture with blue ancestry.

**Figure 3:**
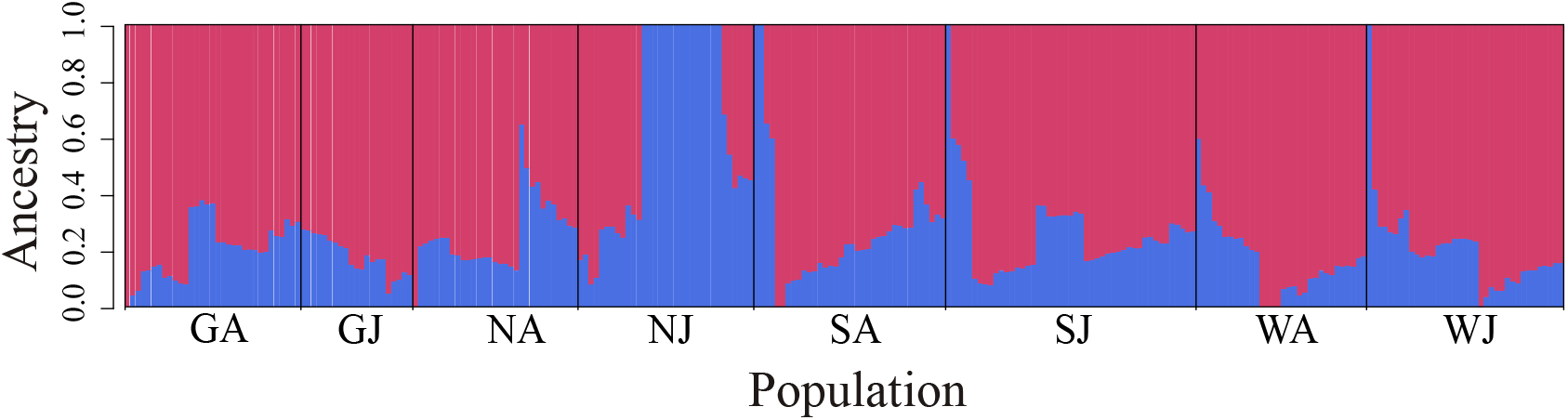
Admixture analysis showing separation of individuals into two genetic clusters, red and blue. Each vertical bar represents a separate individual where the proportion of red and blue indicate the percent ancestry or assignment to each genetic cluster. GA – Goofnuw Adults, GJ – Goofnuw juveniles, NA – Nimpal adults, NJ – Nimpal Juveniles, SA – South Tip adults, SJ – South Tip juveniles, WA – West Outer Reef adults, WJ – West Outer Reef juveniles.

Weighted *F*_ST_ estimates between all combinations of cohorts and sites demonstrates comparable levels of genetic differentiation among groups (supp. table 2). South Tip juveniles have the lowest average *F*_ST_ (0.0366) compared to all other groups, followed by West Outer Reef juveniles (0.0382). Goofnuw Channel juveniles and Nimpal adults had the highest average values of *F*_ST_ (0.0450). *F*_ST_ between exclusively juvenile comparisons was not significantly different compared to *F*_ST_ between exclusively adult comparisons, demonstrating no significant effect of age structure on genetic differentiation (supp. fig. 5).

### Genetic Relatedness and Inbreeding among Nimpal Juveniles

Hierarchical clustering based on genetic distances between samples demonstrated the presence of three clonal pairs in Goofnuw Channel (fig. 4).

**Figure 4:**
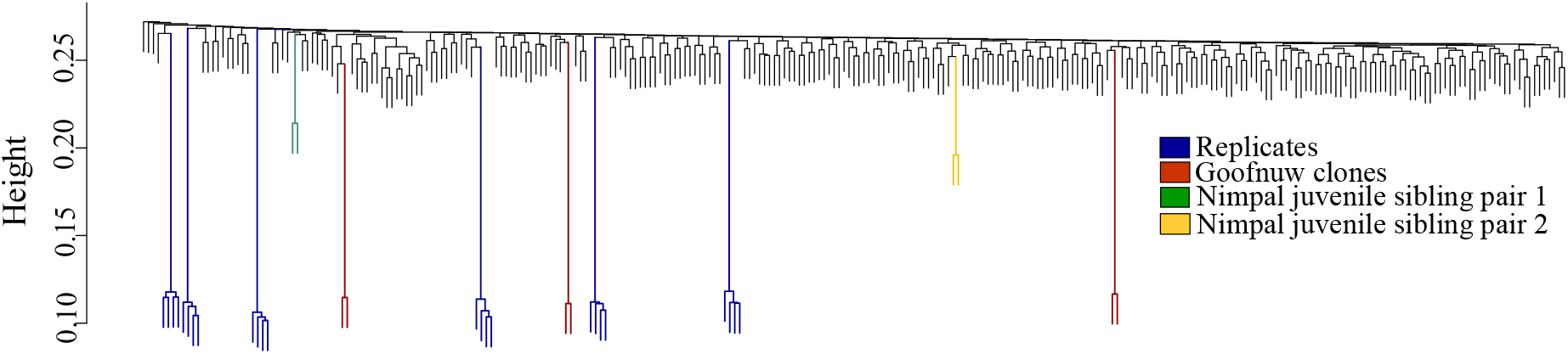
Hierarchical clustering dendrogram of genetic similarity for all individuals. Each individual is represented by a single terminal branch. Blue branches represent technical replicate samples, red branches represent Goofnuw Channel *A. hyacinthus* clones, and the green and yellow branches represent putative sibling pairs of juveniles at Nimpal Reef.

These clones form branches of the same length as our replicate control samples, demonstrating nearly 100% genetic identity between clonal pairs sharing the same branch (fig. 4). Unexpectedly, we found two potential sibling pairs among the juveniles at Nimpal Reef, based on their similarity shown in the hierarchical clustering tree (fig. 4). To corroborate whether these individuals were in fact related, we calculated relatedness values between all combinations of individuals in our dataset using ngsRelate. Our results demonstrate that the juvenile sibling pairs identified in the dendrogram have pairwise relatedness values (0.352 and 0.577) consistent with those found in first degree relatives, most likely siblings or half-siblings (table 1). We also found another two pairs of Nimpal juveniles that showed high pairwise relatedness values (0.110 and 0.103) relative to the entirety of the Yap *A. hyacinthus* population and could potentially be cousins or second cousins (table 1).

**Table 1:**
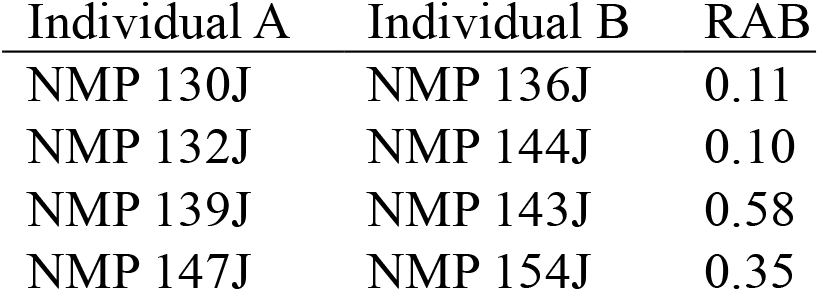
Pairs of individuals with pairwise relatedness values greater than 0.1. RAB indicates the pairwise relatedness value.

| Individual A | Individual B | RAB |
| --- | --- | --- |
| NMP 130J | NMP 136J | 0.11 |
| NMP 132J | NMP 144J | 0.10 |
| NMP 139J | NMP 143J | 0.58 |
| NMP 147J | NMP 154J | 0.35 |

Since discovery of genetically related individuals at the same site is very unexpected in a broadcast-spawning coral species, we sought to determine whether Nimpal juveniles in general had higher levels of intracohort relatedness compared to the other cohorts on Yap Island. Although the range of relatedness values is quite large, Nimpal juveniles do have higher overall relatedness to each other compared to relatedness within other cohorts (supp. fig. 6B). To further refine the analysis, all individuals were sorted into quartiles based on their proportion of blue ancestry as shown in the admixture analysis, since Nimpal juveniles comprise the majority of individuals with 100% blue ancestry (fig. 3). Those with a higher proportion of blue ancestry showed higher relatedness values amongst each other (fig. 5B).

**Figure 5:**
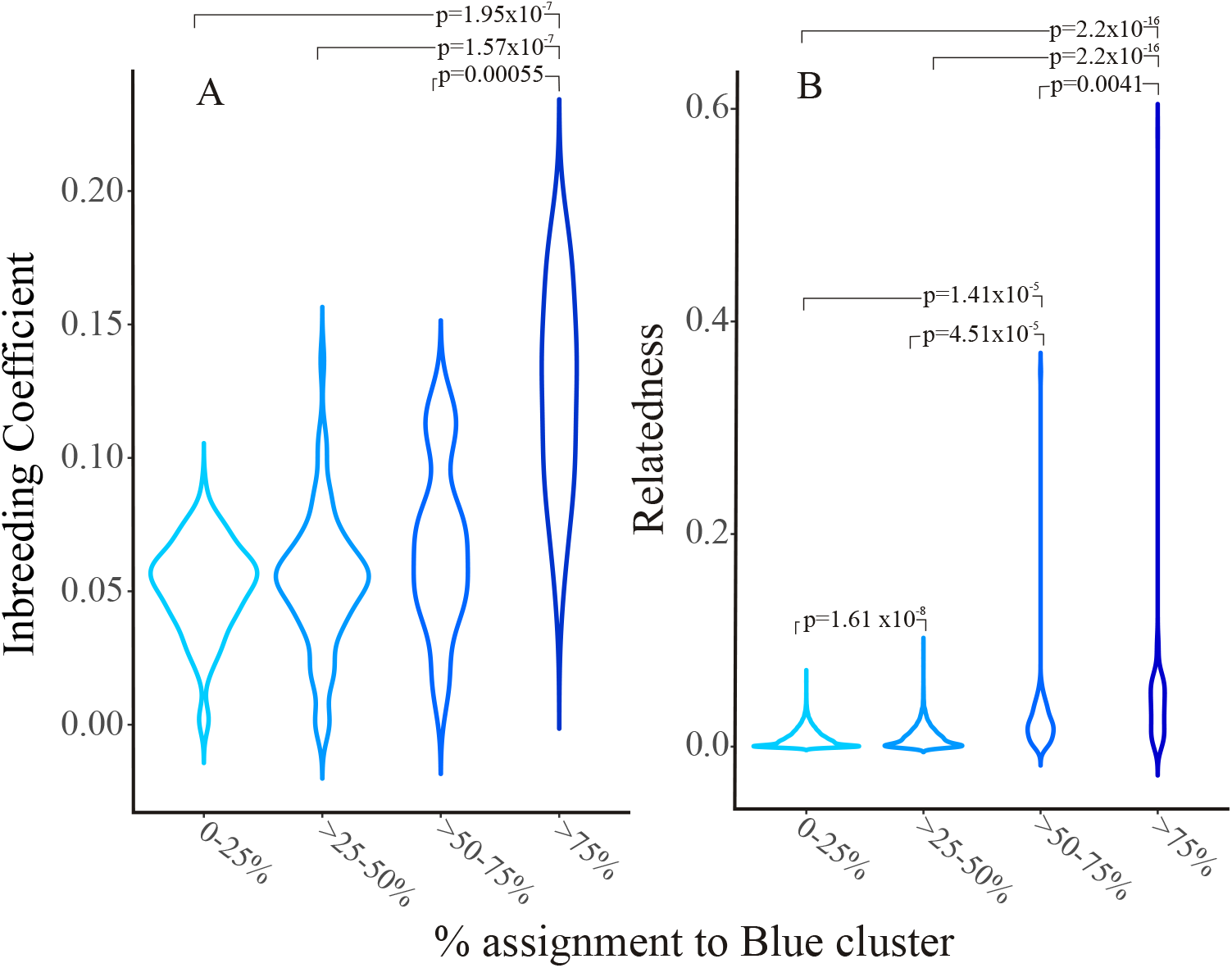
Inbreeding coefficients (A) and pairwise relatedness (B) of all individuals separated into quartiles based on the percent assignment to the blue admixture cluster.

Individuals with greater than 75% assignment to the blue cluster had significantly greater levels of relatedness compared to individuals with 50-75% blue (p=0.0041), 25-50% blue (p=2.2×10^−16^), and 0-25% blue (p=2.2×10^−16^).

If genetically related individuals were settling close to each other for more than one generation, this should result in elevated levels of inbreeding. Indeed, this is what we find in individuals with a large proportion of blue ancestry, comprised mostly of Nimpal juveniles (supp. fig. 6A, fig. 5A). Those individuals with greater than 75% assignment to the blue cluster had significantly greater inbreeding coefficients compared to the 50-75% quartile (p=0.00055), the 25-50% quartile (p=1.57×10^−7^), and the 0-25% quartile (p=1.95×10^−7^). Of the Nimpal juveniles, 15 out of 35 individuals show 100% genetic affiliation with the blue cluster, whereas the others are admixed. The remaining 20 out of 35 Nimpal juveniles and all Nimpal adults belong to the main panmictic population on Yap Island, are characterized by a reduced proportion of blue ancestry, low inbreeding coefficients and minimal levels of relatedness with each other (figs. 3, 5A-B). Similarly, we find significantly lower levels of heterozygosity in Nimpal juveniles with entirely blue ancestry compared to those Nimpal juveniles with mixed ancestry, as is expected when inbreeding levels are elevated (fig. 6).

**Figure 6:**
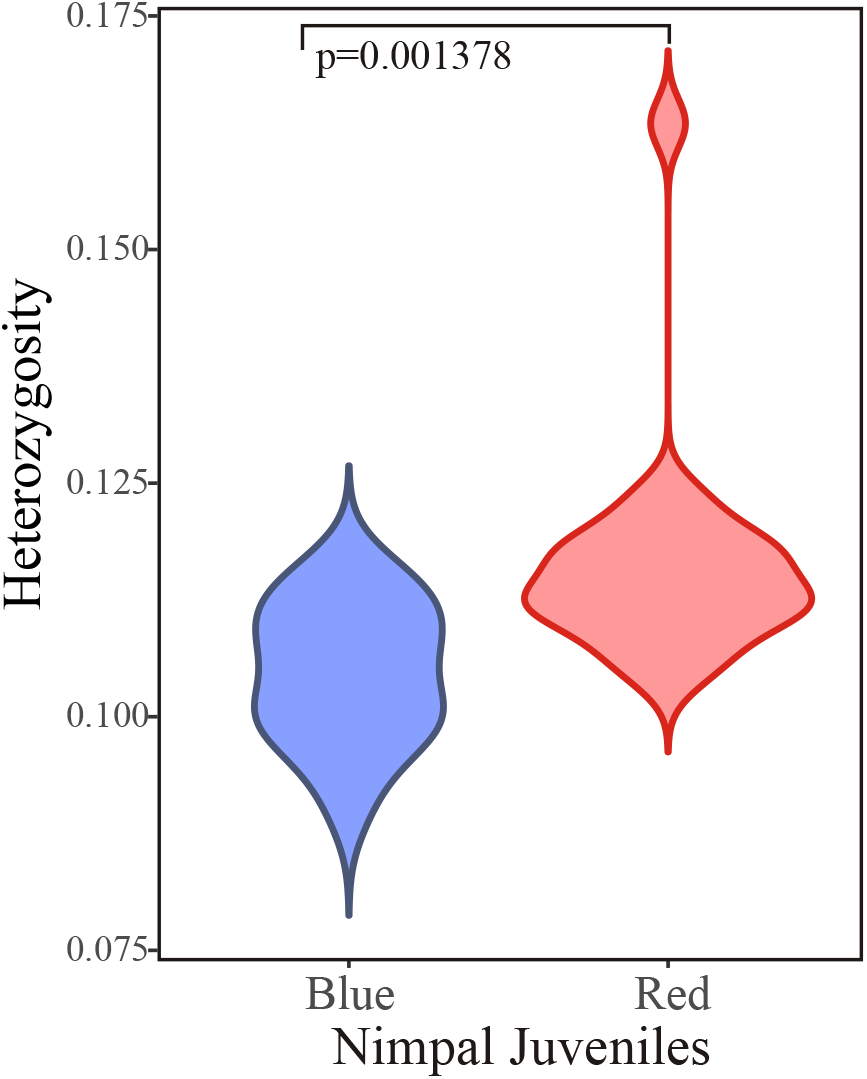
Heterozygosity of Nimpal juveniles with 100% blue ancestry (n=15) compared to red-blue admixed Nimpal juveniles (n=20).

### Genomic signals in inbred Nimpal juveniles

Based on the greater level of inbreeding found in a large proportion of Nimpal juveniles, we expected to find an overall reduction in genetic diversity of this cohort compared to other *A. hyacinthus* cohorts. When comparing theta (genetic diversity) profiles across the genome in Nimpal juveniles from the inbred, blue ancestry group (n=15) to the “panmictic” Nimpal juveniles (n=20), we find overall lower baseline levels of diversity across the genome (fig. 7).

**Figure 7:**
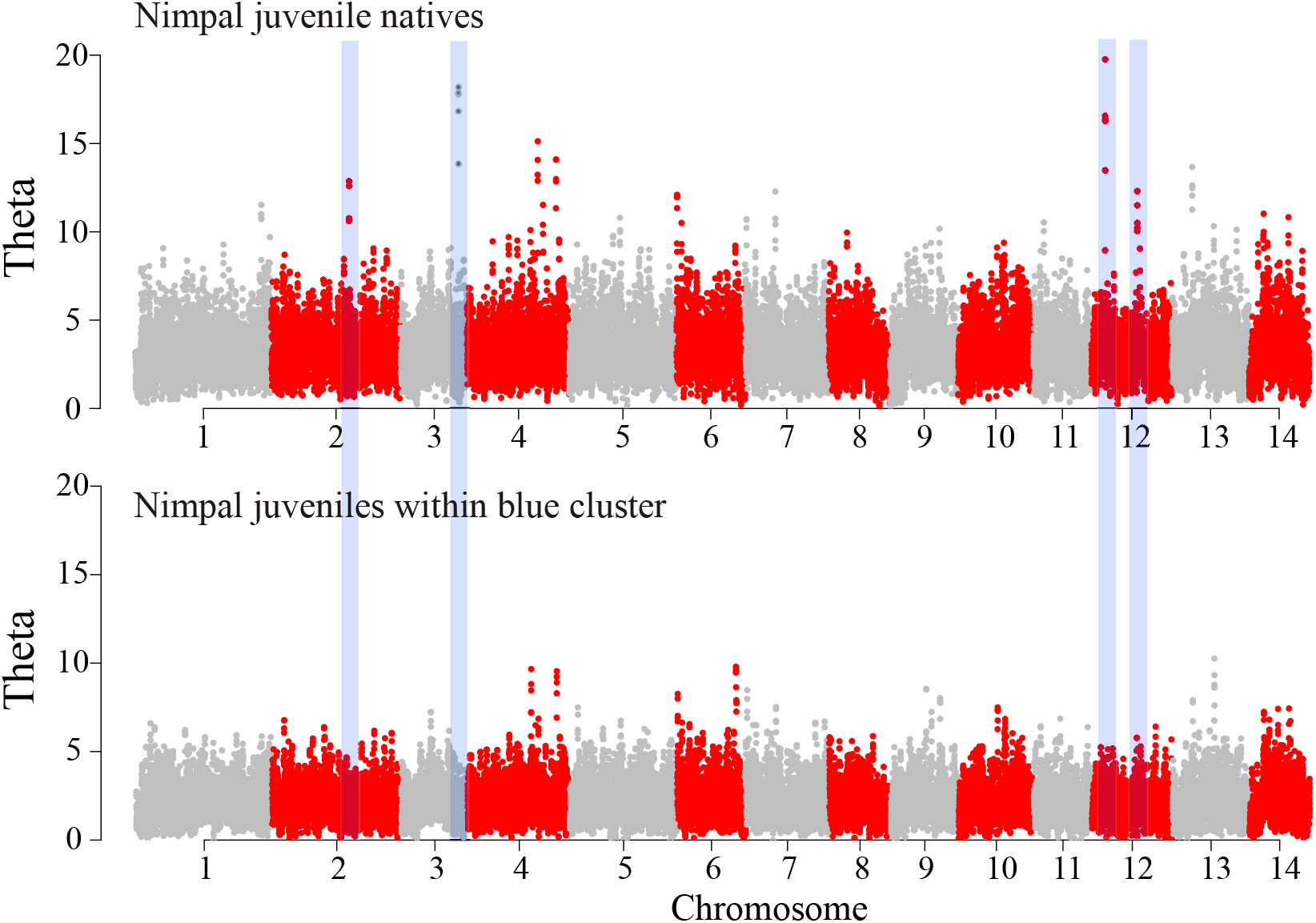
Manhattan plot of theta values across 50kb windows and 10kb step size. Peaks represent regions of high diversity and vice versa. Top manhattan plot shows values for Nimpal juvenile natives (i.e. admixed individuals) (n=20) and the bottom manhattan plot shows theta values for the Nimpal juveniles with 100% blue ancestry (n=15).

On average, the blue ancestry Nimpal juveniles are half as genetically diverse across the genome compared to all other cohorts. Additionally, there are a few regions (middle of chromosome 2, end of chromosome 3, and two segments within chromosome 12) that are highly polymorphic among individuals representing the panmictic population that show dramatic reductions in diversity in the blue ancestry Nimpal juveniles (fig. 7 and supp. fig. 7).

### Sequence coverage

Since it is possible that our observations of higher inbreeding and relatedness and decreased heterozygosity in the blue ancestry group could in part be due to lower sequence coverage, we performed a principal component analysis to determine if any of genetic measures (inbreeding, heterozygosity, percent blue ancestry) were driven by sequencing coverage (log_10_ read depth). We find that, inbreeding, percent blue ancestry, and heterozygosity load in the direction of PC1, whereas read depth is loaded in the orthogonal direction of PC2 (supp. fig. 8), which indicates that variation in read depth is not a major driver of variation in genetic measures.

As a corollary analysis, we looked at where the relatives (siblings, cousins) fell within the distribution of sequence coverages (supp. fig. 9). One cousin pair was at the lower end of the distribution of sequence coverages, and thus, it is possible that the estimates of relatedness are slightly inflated for this pair of relatives. However, the other cousin pair and both sibling pairs fall well within the range of adequate sequence coverage for accurate genotyping. The log_10_ read depth was not significantly different between Nimpal juveniles belonging to the blue cluster (n=15) and the other Nimpal juveniles (n=20) (supp. fig. 10).

## Discussion

### Intergenerational stability in genetic sources of recruits among A. hyacinthus cohorts

Among the four sites sampled, juvenile cohorts from Goofnuw Channel, South Tip, and West Outer Reef all demonstrate evidence for stability in genetic composition compared to the adult population. There is no evidence for intergenerational shifts in allele frequencies or genetic diversity, as evidence by PCoA and genome-wide theta values (fig. 2A-D, fig. 7., supp. fig. 7). Furthermore, at South Tip there was a 6-year gap between the sampling of the adult and juvenile cohorts, due to a crown-of-thorns outbreak that extirpated most of the adult acroporid colonies about a year after the initial adult collection. Still, the genetic composition of newly recruited South Tip juveniles was remarkably similar to the pre-disturbance adult cohort, demonstrating both stability in the genetic sources of recruits and the potential for rapid recovery of local coral populations after disturbance. Indeed, we find that genetic diversity in juveniles at South Tip is similar to the extirpated adult population, while pairwise *F*_ST_ between them and other cohorts is slightly lower compared to other cohort comparisons (average F_ST_ of 0.0363 vs 0.0418) (supp. table 2). Additionally, our PCoA shows extensive overlap between the juveniles and former adults at South Tip (fig. 2), and our admixture analysis indicates extensive genetic similarity between South Tip juveniles and the majority of the Yap *A. hyacinthus* population (fig. 3). Together, these results indicate that the recruit pool is genetically well-mixed and representative of the whole panmictic *A. hyacinthus* population on Yap Island, which has allowed for recovery of genetic diversity at South Tip in a single generation.

Ensuring successful recruitment of larvae to these reefs is vital, since most *A. hyacinthus* colonies are likely the product of sexual versus asexual reproduction. Only three pairs of asexual clones were identified among all the sampled *A. hyacinthus* colonies, and these were from Goofnuw Channel Reef, which is on the eastern side of Yap Island and is subject to greater storm damage. Considering that *A. hyacintuus* is a brittle, table-forming coral species, the contribution of asexual reproduction to the generation of new colonies is lower than we expected. However, there was a healthy abundance of acroporid and non-acroporid adult and juvenile colonies at all sites, except Goofnuw, suggesting Yap Island reefs have robust levels of recruitment and an age-structure suitable for continued maintenance of coral populations (supp. fig. 4).

### Co-recruitment of related larvae leads to elevated inbreeding and relatedness

Larvae of *Acropora hyacinthus* spend several days to weeks drifting with ocean currents before settling on the reef (Shanks, Grantham, & Carr, 2003). Finding not just one, but two pairs of sibling or half-sibling juveniles at Nimpal Reef, as well as two pairs of potential cousins or second cousin juveniles (fig. 4, table 1), is therefore entirely unexpected. Although the exact relationship among these corals (sibling vs half sibling and cousins vs second cousins) cannot be definitively established due to generally higher inbreeding in this group, this level of relatedness among sympatric individuals has, to our knowledge, never been reported in any broadcast-spawning coral. Because Nimpal juveniles showing elevated relatedness also show elevated inbreeding (fig. 5A-B), this indicates that their parents have been cross-breeding with relatives, which in turn implies that they have also settled near relatives. Proximity between individuals is one of the most significant factors affecting the potential to cross-fertilize in corals (Yund, 2000). Therefore, co-recruitment, the dispersal and settlement of a genetically related larval cohort to the same reef, has most likely occurred over at least two generations in this genetic lineage. This greatly contrasts with the other *A. hyacinthus* cohorts (South Tip, Goofnuw, West Outer), which exhibit panmixia.

While it is possible that dramatically reduced planktonic larval duration and/or re-circulation of currents near the reef could have led to minimal dispersal of offspring throughout the water column, we would likely have found offspring settled near and amongst their parents; yet we did not find any adults of 100% blue ancestry at Nimpal Reef or at the nearby West Outer Reef. While it is still possible that there is a small enclave of “blue” adults in the area that we did not sample, reduced planktonic larval duration alone seems insufficient to explain co-recruitment of relatives away from their parents. Still, limited dispersal is one potential explanation that cannot be completely ruled out for the emergence of this group.

Another mechanism that can bring genetically related coral larvae together on the same reef is collective or cohesive dispersal of larvae from the same cohort. The abundance of “blue” Nimpal juveniles compared to the number of progenitor adults could be the result of a cohesively dispersed cohort constituting the majority of recruits during one reproductive season, also known as Sweepstakes Reproductive Success (SRS) (Hedgecock & Pudovkin, 2011). However, this would have to occur over multiple generations to result in inbreeding. Still, we may have detected a sweepstakes recruitment event from some highly isolated reef harboring a small and mostly self-seeding, and therefore inbred, *A. hyacinthus* population, although such is the less parsimonious explanation. Generally, evidence for extreme sweepstakes reproduction events, as is suggested by our data, is rare among marine spawners (Flowers, Schroeter, & Burton, 2002; Levitan, 2005), although SRS is compatible with a collective larval dispersal scenario.

Collective dispersal via sibling aggregations has been found in various fishes, but the mechanism of dispersal is thought to be aided by behavioral, auditory, and/or olfactory cues (Christie, Johnson, Stallings, & Hixon, 2010; Kingsford et al., 2002; Pusack, Christie, Johnson, Stallings, & Hixon, 2014). In contrast, evidence for cohesive dispersal in marine invertebrates is still rare. Riquet et al. (2017), however, find co-settled full and half-siblings of the brooding gastropod *Crepidula fornicata* over the course of three years, suggesting collective dispersal of larvae occurs in this invertebrate (Riquet, Comtet, Broquet, & Viard, 2017). Moreover, they also identify inbred larvae among their samples, analogous to what we have found in the Nimpal juveniles. Similarly, Iacchei et al. (2013) find that populations of California spiny lobster exhibit a patchy genetic structure with a majority of sites demonstrating significantly greater kinship levels than expected by chance, again suggesting that larvae maintain planktonic cohesiveness during dispersal (Iacchei et al., 2013). While it seems highly unlikely that related coral larvae would stay together in the plankton via sensory mechanisms, a passive mechanism such as a mucus net produced by parents during spawning is not inconceivable. Coral mucus helps to keep eggs and sperm from dispersing widely to increase fertilization success, and thus might also keep larvae aggregated (Oliver & Willis, 1987). In addition, oceanographic models of larval dispersal demonstrate stochasticity and show that larvae do not mix in the water column as much as previously thought, increasing the prospect that cohesive larval dispersal could be facilitated by currents (Siegel et al., 2008). Knowledge of the natural behavior of coral larvae in the ocean is also very limited, leaving open the possibility for unexpected dispersal patterns.

### Potential incipient cryptic species formation event in Yap A. hyacinthus *corals*

Our genome-wide scan of theta values revealed the presence of four regions of greatly reduced diversity in the blue ancestry Nimpal juveniles that are highly diverse in all other cohorts (fig. 7, supp. table 3). A total of ten genes lie directly within these regions, including three unannotated genes, a notch signaling ligand (DLL4), XPO6, PKDIL2, WDR87, SNRPE, LRMP, and NME5. Out of the seven annotated genes, four (LRMP, PKDIL2, WDR87, and NME5) have putative functions related to fertilization or have closely related paralogs involved in fertilization and/or spermiogenesis (supp. table 3). A close paralog of PKDIL2 (PKDREJ) generates a Ca^2+^ transporting channel involved in initiating the acrosome reaction of sperm and is a candidate sperm-receptor gene in humans (Hamm, Mautz, Wolfner, Aquadro, & Swanson, 2007; Sutton, Jungnickel, Ward, Harris, & Florman, 2006). The paralog of WDR87 is WDR25, which is an epididymis secretory sperm binding protein. The sea urchin ortholog of NME5 (NME8) is implicated in sperm ciliary function, and this protein may be required for sperm tail maturation (Munier et al., 2003). NME5 has also been suggested to function in spermatogenesis through the elimination of reactive oxygen species (Choi et al., 2009; McReynolds et al., 2014). LRMP is thought to play a role in fertilization during pronucleus congression and fusion (Nguyen, Ishihara, Wuhr, & Mitchison, 2012). In addition, approximately 50kb downstream of the high diversity region on chromosome 3 are two genes, one of which (CSE1L) is annotated as an epididymis secretory sperm binding protein and is highly expressed in human placenta, ovaries, and testis. Although the number of observations is too limited to show a differential enrichment signal, the high proportion of genes with functions involved in fertilization in regions of reduced diversity suggests that modification of mating compatibility within the blue ancestry group could be one of the consequences.

The loss of genetic diversity at these loci could have been driven by directional selection to improve fertilization among relatives, if sperm is limited for example (Levitan & Petersen, 1995). Alternatively, loss of genetic diversity in these loci could have resulted from relaxation of diversifying selection if these loci function to limit polyspermy (Levitan, terHorst, & Fogarty, 2007). In any case, this diversity loss may have resulted in at least partial incompatibility of the blue cluster with the rest of the population. This hypothesis is the more likely considering that at least some reproductive isolation would be necessary to prevent this group from being genetically swamped via outbreeding with native genotypes, since *A. hyacinthus* is abundant on Yap Island. Additionally, there is a greater probability of generating offspring with elevated inbreeding and relatedness if mating is restricted within a subset of adults. Co-recruitment could also facilitate the propagation of a partially or fully reproductively isolated group, as it should increase mating success due to co-localization of compatible mates.

In broadcast spawning marine invertebrates such as corals, there are few cues to limit mating between species, besides spawning time and sperm-egg interaction proteins. Thus, in theory, changes in only a handful of fertilization-related genes could be sufficient to generate reproductively isolated coral species (Evans & Sherman, 2013; Hart, Sunday, Popovic, Learning, & Konrad, 2014; Levitan et al., 2004; Palumbi, 1994; Palumbi, 2009). It is also telling that we find very few individuals with a proportion of blue ancestry greater than 50% but less than 100%, suggesting that backcrosses of red/blue hybrids to individuals of pure blue ancestry are rare (supp. table 4), perhaps as a result of an incomplete reproductive barrier.

Ladner and Palumbi (2014) show that *A, hyacinthus* is part of a cryptic species complex in the Indo-Pacific that includes at least four species with extensive sympatry and evidence for introgression (Ladner & Palumbi, 2012). In Palau, the next closest reef to Micronesia that contains *A. hyacinthus*, Ladner and Palumbi (2012) detected three of these cryptic *A. hyacinthus* species. The evidence presented here suggests that another incipient cryptic speciation event might be underway at Yap Island, although the evolutionary persistence of this group is uncertain. From our photos, we could find no obvious morphological differences between inbred and panmictic Nimpal juveniles. The relative isolation of Yap Island from other coral sources could be a prime factor contributing to the emergence of this new genetic subdivision. For example, Budd and Pandolfi (2010) demonstrate a tendency for Caribbean Pleistocene corals at the edge of their distributions to split into different lineages owing to reduced gene flow to these regions (Budd & Pandolfi, 2010), and Richards et al. (2016) highlight evidence for the existence of several cryptic acroporid species separated across large oceanic distances, which were previously thought to be panmictic (Richards, Berry, & Van Oppen, 2016).

Our findings highlight previously underappreciated potential for recruitment processes to influence genetic subdivisions in a broadcast spawning coral. Taken together, our results suggest that co-recruitment of genetically related individuals could be contributing to the emergence and establishment of a new cryptic *A. hyacinthus* species, or at least a genetically distinct subpopulation.

## Acknowledgements

We thank the Yap Community Action program (Yap CAP) and Dr. Rong Ma for assisting with fieldwork planning and sample collection, and Ka’ohinani Kawahigashi for help with molecular work.

## Authors’ contributions

S.B, S.W.D., and M.V.M. conceived of the study. S.W.D. contributed *A. hyacinthus* samples. S.J.B. designed the study, conducted sampling and bioinformatic analyses, performed molecular work and wrote the first draft of the paper. S.B., S.W.D., and M.V.M. contributed to revisions.

## Data Accessibility

All sequencing data generated from the project are available from NCIB-SRA, accession no. PRJNA565239

