## Supplemental Table 2 for "Co-recruitment of relatives in a broadcast-spawning coral (*Acropora hyacinthus*) facilitates emergence of an inbred, genetically distinct group within a panmictic population"

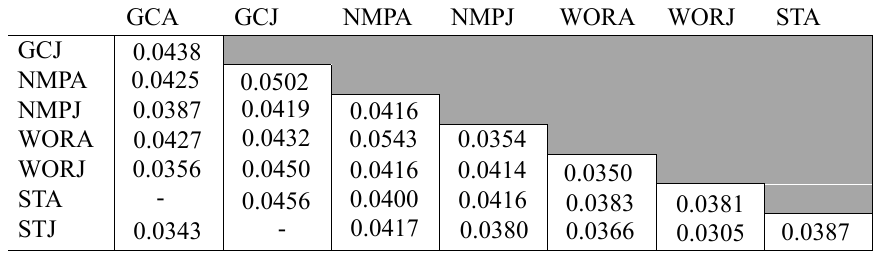

Supplemental Table 2: Pairwise *F*_ST_ values for all population cohort comparisons. For population labels, GC indicates Goofnuw Channel, NMP indicates Nimpal, WOR indicates West Outer Reef, and ST indicates South Tip. Cohort labels are “A”, indicating adults and “J”, indicating juveniles.
